# Trophic ecology of deep-sea megafauna in the ultra-oligotrophic Southeastern Mediterranean Sea

**DOI:** 10.1101/2022.01.20.477062

**Authors:** Tamar Guy-Haim, Nir Stern, Guy Sisma-Ventura

## Abstract

The trophic ecology of fourteen species of bathybenthic and bathypelagic fishes and six species of bathybenthic decapod crustaceans from the continental slope and rise of the Southeastern Mediterranean Sea (SEMS) was examined using stable isotope analysis. Mean δ^13^C values among fish species varied by more than 4.0‰, from −20.85‰ (*Macroramphosus scolopax*) to −16.57‰ (*Conger conger* and *Centrophorus granulosus*), and increased as a function of depth (200 - 1400 m). Mean δ^13^C values of the crustaceans showed smaller variation, between −16.38‰ (*Polycheles typhlops*) and −18.50‰ (*Aristeus antennatus*). This suggests a shift from pelagic to regenerated benthic carbon sources with depth. Benthic carbon regeneration is further supported by the low benthic-pelagic POM-δ^13^C values, averaging −24.7 ± 1.2‰, and the mixing model results, presenting very low contribution of epipelagic POM to the bathyal fauna. Mean δ^15^N values of fish and crustacean species ranged 7.91 ± 0.36‰ to 11.36 ± 0.39‰ and 6.15 ± 0.31‰ to 7.69 ± 0.37‰, respectively, resulting in trophic position estimates, occupying the third and the fourth trophic levels. Thus, despite the proximity to the more productive areas of the shallow shelf, low number of trophic levels (TL~1.0) and narrow isotopic niche breadths (SEAc <1) were observed for bathybenthic crustaceans (TL = 3.62 ± 0.22) and bathypelagic fishes (TL = 4.33 ± 0.34) in the study area – probably due to the ultra-oligotrophic state of the SEMS resulting in limited carbon sources. Our results, which provide the first trophic description of deep-sea megafauna in the SEMS, offer insight into the carbon sources and food web structure of deep-sea ecosystems in oligotrophic marginal seas, and can be further used in ecological modeling and support the sustainable management of marine resources in the deep Levantine Sea.

## 1 Introduction

Deep-sea ecosystems cover much of the oceans seafloor and play a major role in large-scale biogeochemical cycles (Walsh, 1991;Drazen and Sutton, 2017). They provide ecosystem services that are important to humans, including carbon sequestration, nutrient recycling and burial, waste accumulation and fisheries production (Danovaro et al., 2008;Mengerink et al., 2014;Thurber et al., 2014). Recent studies have shown that an increasing number of stressors, including climate change (warming), deoxygenation, ocean acidification, as well as, overfishing, and natural resource extraction (e.g., Stramma et al., 2008;Yasuhara et al., 2008;Stramma et al., 2010;Helm et al., 2011;Tecchio et al., 2015) are expanding into deep environments, thus threatening the diversity and stability of deep-sea ecosystems. Consequently, studying the status of deep-sea communities and describing deep-sea ecosystem structures are currently gaining more and more attention.

Continental slopes account for ~11% of the total ocean floor (Ramirez-Llodra et al., 2010), connecting the shallow shelf productive areas with the abyssal plains along steep seabed gradients. Covering large bathymetric ranges (~ 200 – 2000 m), these dynamic habitats exhibit strong spatial differences in temperature, salinity, nutrient concentrations and consequently, in habitat suitability (Koslow, 1993;Gordon et al., 1995;Neat et al., 2008;Bergstad, 2013;Pajuelo et al., 2016). Although bathyal habitats are relatively isolated from terrestrial inputs, they can support diverse deep-sea fauna (Gordon and Swan, 1997;Kelly et al., 1998;Menezes et al., 2006;Neat et al., 2008), even in ultra-oligotrophic basins, such as the easternmost Mediterranean Sea (Goren et al., 2008). In deep-sea benthic ecosystems, fish can play key ecological and biogeochemical roles (Drazen and Sutton, 2017) by regulating nutrient limitation and zooplankton populations (Hopkins and Gartner, 1992;Pakhomov et al., 1996).

Deep-benthic ecosystems largely rely on particulate organic matter (POM) that passively sinks from the surface waters or by lateral transport as a primary source of nutrients (Tecchio et al., 2013). Animals that carry out vertical diel migrations through the water column (Trueman et al., 2014) and occasional sink of large animal carcasses is another important food source to deep ecosystems (Smith and Baco, 2003). Each of these primary food sources carry a distinct isotopic signature that may reflect its origin, resulting from different chemo-physical processes. Thus, by knowing the isotopic composition of the food source that fuels a specific food web, it is possible to reconstruct the trophic structure and dynamics of specific habitats (Post, 2002).

Stable isotope analysis (SIA) has been used successfully to study trophic level, important prey types, and trophic niche breadth in deep-sea ecosystems (e.g., Boyle et al., 2012;Shipley et al., 2017a). Nitrogen stable-isotope composition (δ^15^N) is used to determine the trophic position of an animal, as it preferentially fractionates as a function of its diet, where the heavy isotopes are retained in the consumers in respect to their prey by 2–4‰ (Post, 2002). Carbon isotopes (δ^13^C) fractionate much less with each trophic step (<1‰), but can be effectively used to infer basal sources of carbon. Moreover, SIA provides an integrated view of an organism’s diet over time-scales relevant to tissue turnover rates rather than digestion rates (Peterson and Fry, 1987;Post, 2002), thereby providing estimates of the trophic position of an organism within a specific food web.

Knowledge of food web structure and dynamics is key to our understanding of ecological communities and their functioning (Polis and Strong, 1996;Winemiller and Polis, 1996). This fundamental information is, however, lacking in many oceanographic regions, including the Southeastern Mediterranean Sea (hereafter, SEMS) (Parzanini et al., 2019) – one of the most oligotrophic, nutrient-impoverished marginal basin, worldwide (Kress et al., 2014). The SEMS provides a miniature model of processes occurring in vast oligotrophic gyres, an ideal location to study food web structure and functioning under sever nutrient limitation. Furthermore, the SEMS is one of the regions where sea surface temperatures are rising at the fastest rates under recent climate changes (Sisma-Ventura et al., 2014; Ozer et al., 2017) and is one of most vulnerable marine regions to species invasions (Rilov and Galil, 2009), which have been also reported from deep-sea habitats (Galil et al., 2019). Understanding deep-sea community structure and functioning is of prime importance for developing better predictions regarding the ecological effects of future climate change.

To date, much of the research describing the trophic ecology of the Eastern Mediterranean Sea (EMS) has focused on zooplankton groups (Koppelmann et al., 2003;Koppelmann et al., 2009;Hannides et al., 2015;Protopapa et al., 2019), shallow rocky reefs (Fanelli et al., 2015), and on anthropogenically-influenced coastal environments (Grossowicz et al., 2019), while less attention has been paid to deep-sea fishes and crustaceans that occupy higher trophic levels. Here we used bulk carbon and nitrogen stable isotopes (δ^13^C and δ^15^N) to study the trophic ecology of bathypelagic and bathybenthic fishes and crustaceans from the southeast Mediterranean continental slope and rise. We explored potential factors that may explain the variability in isotope values across species. These data offer insights into the carbon sources and trophic complexity of deep-sea ecosystems in oligotrophic marginal seas.

## 2 Materials and methods

### 2.1 Study sites and sampling design

Sampling campaigns were conducted in the course of three oceanographic cruises during 2017-2019, as part of the national deep-water monitoring program of the Israeli Mediterranean Sea performed by Israel Oceanographic and Limnological Research (IOLR). Sampling sites were divided to three major benthic habitats: (1) the end of the continental shelf, with an average depth of 200 m; (2) the continental slope with depth range of 500-600 m; and (3) the deep bathyal plateau (continental rise) with depth range of 1000-1400 m (**Figure 1**). Specimens were collected onboard the R/V *Bat-Galim*, using a semi-balloon trawl net with an opening of eight meters and mesh size of 10 mm. Once the trawls were retrieved, animals were sorted, enumerated, weighted and visually identified to species level. The total length (cm) of each specimen was recorded. Specimens for SIA were selected and frozen whole at −20 °C until processed at the IOLR.

**Figure 1.**
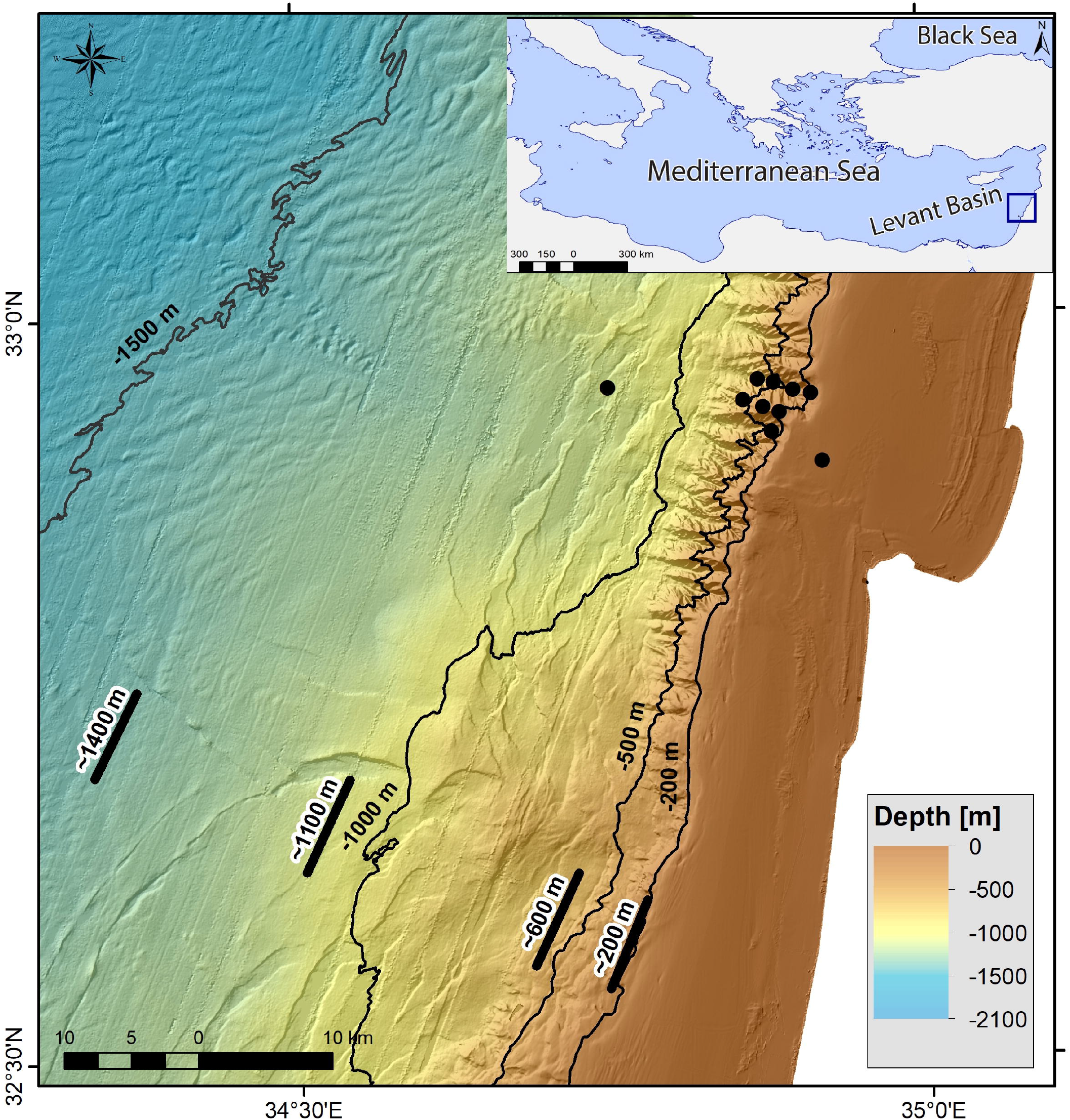
Map of sampling sites in the SEMS. The locations of POM samples are presented in black circles, and locations of fishes and crustaceans by trawl net are presented in black lines.

POM was collected during three research expeditions in winter 2018, summer 2020, and winter 2021, across the shelf, slope and rise of the SEMS (**Figure 1**). POM samples were collected throughout the water column using 8-L Niskin bottles. Water samples were then filtered on pre-combusted 47-mm glass fiber filters (Whatman) in duplicates at low pressure and dried at 60 °C for 24 h prior to isotope analysis.

### 2.2 Stable Isotopes analysis

SIA was conducted on 86 fish and 46 crustacean specimens as well as 77 POM samples (**Table 1**). White muscle tissue for SIA was dissected from the dorsal musculature of fishes and from the abdominal segment of the crustaceans. Samples were rinsed with deionized water, frozen, and lyophilized for 48 h. Freeze-dried samples were homogenized using a mortar and pestle, weighed, and shipped to the Stable Isotope Facility at Cornell University for SIA analysis. The isotopic composition of organic carbon and nitrogen was determined by the analysis of CO_2_ and N_2_ continuous-flow produced by combustion on a Carlo Erba NC2500 connected on-line to a DeltaV isotope ratio mass spectrometer coupled with a ConFlo III interface.

**Table 1:**
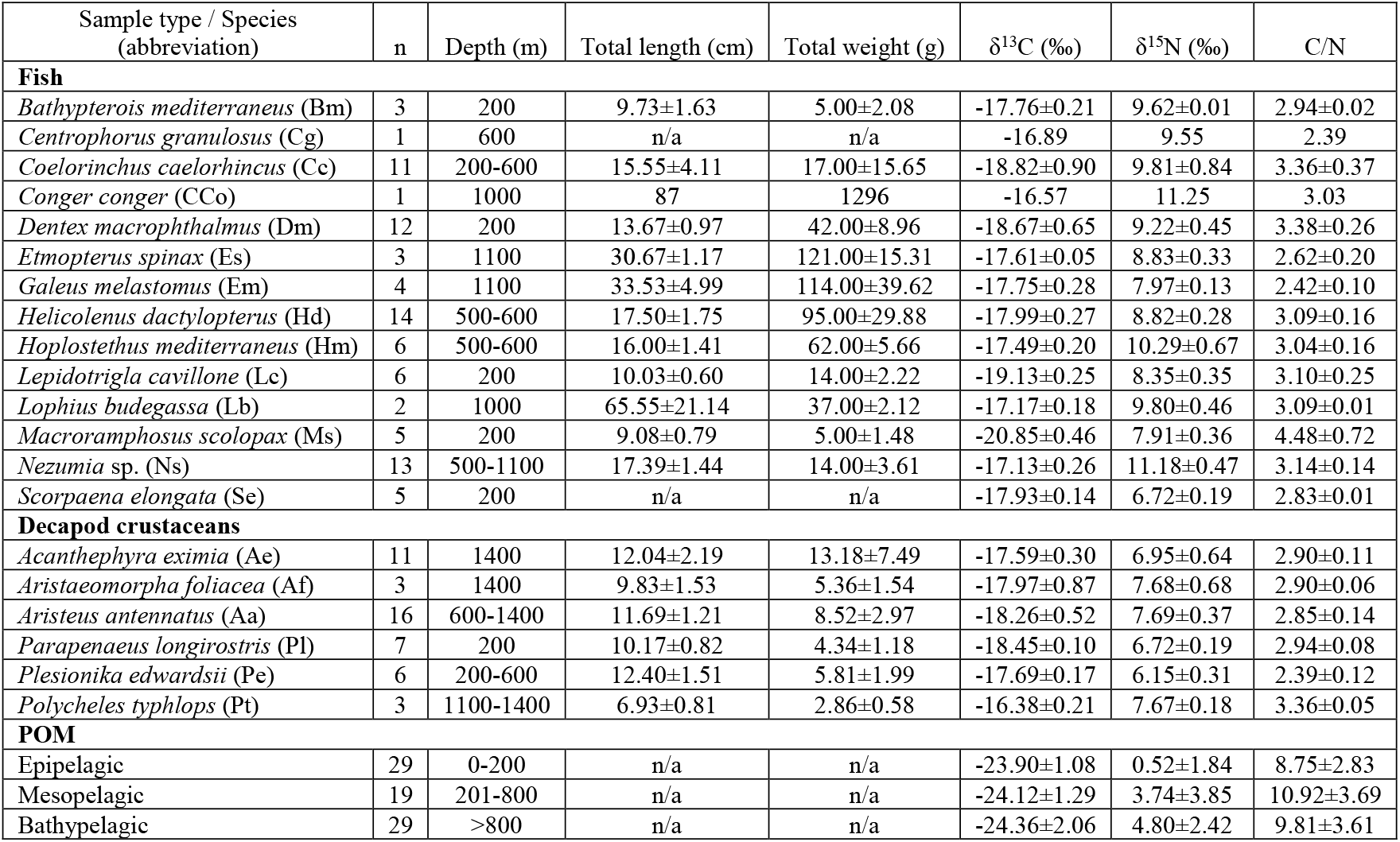
Sample descriptions and bulk δ^13^C and δ^15^N isotope data (mean ± SD).

Measured isotope ratios are reported in the δ-notation, i.e., as the deviation in per mill (‰) from the international standards:

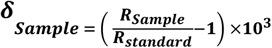

where, R represents the ^15^N/^14^N or ^13^C/^12^C ratio. Stable isotope data are expressed relative to international standards of Vienna PeeDee belemnite and atmospheric N_2_ for carbon and nitrogen, respectively. The analytical precision for the in-house standard was ± 0.04‰ [1σ] for both δ^13^C and δ^15^N. The C/N ratios of fishes and crustaceans in this study were low (species mean C/N ranged between 2.33–4.48; where in 97% of individuals C/N < 4.0, see **Supplementary Figures 1, 2**), suggesting that lipids did not significantly affect the δ^13^C interpretation (Post et al., 2007). Therefore, all data analyses were performed on uncorrected δ^13^C values. To determine if the isotopic signatures of POM samples changed with depth, we used collection depth to classify POM samples as epipelagic (0–200 m), mesopelagic (200–800 m), or bathypelagic (>800 m).

### 2.3 Data analysis

The trophic position (Trophic Level, TrL_i_) was calculated for each species according to Post et al. (2007):

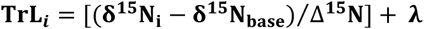

where, δ^15^*N_i_* is the mean species δ^15^N, and δ^15^*N_base_* stands for the primary producer or primary consumer being used to set the isotopic baseline. We applied the trophic discrimination factor Δ^15^*N* of 3.15‰, which was previously used to calculate the trophic level of meso- and bathypelagic fish (Valls et al., 2014;Richards et al., 2018). The *λ* represents the trophic level of the organism being used to set the baseline. Following Protopapa et al. (2019), epipelagic POM was set as the baseline and *λ* was set to an intermediate value of 1.5, since it consists mostly of phytoplankton (TL = 1) and micro- and mesozooplankton (TL = 2) (Albo-Puigserver et al., 2016), equally contributing due to intensive top-down control in this region (Belkin et al., 2022).

Least-squares linear regression analysis was conducted for each species to explore the relationship between fish length and the δ^13^C and δ^15^N values. Spatial variation in δ^13^C and δ^15^N of both fishes and crustaceans was investigated using least-squares linear regression between isotopic values and bathymetric depths. All statistical analyses were performed in R v. 4.0.5 (R Core Team, 2020).

The trophic breadth of each species (*n* >3) and trophic similarity among species were assessed by calculating Standard Ellipse Area (SEA) using the R package SIBER v. 2.1.6 (Jackson et al., 2011;Jackson and Parnell, 2021). Size-corrected SEAs (SEAc) were calculated for each species, which adjusts for underestimation of ellipse area at small sample sizes and allows for inter-study comparison of ellipse sizes (Jackson et al., 2011). Fish and crustacean community metrics were calculated based on Layman et al. (2007).

Bayesian mixing models were applied using R package MixSIAR v. 3.1.12 (Stock et al., 2018;Stock et al., 2021) to estimate the relative contribution of epi-, meso-, and bathypelagic POM to each species. These models are sensitive to variable discrimination factors (Bond and Diamond, 2011;Olin et al., 2013), which may be influenced by diet (Caut et al., 2009), tissue type (Malpica-Cruz et al., 2012), temperature (Britton and Busst, 2018), and species-specific metabolic rates (Pecquerie et al., 2010). Largely, greater δ^15^N discrimination factors (>3.0‰) are associated with lower trophic-level species, and are significantly lower (<3.0‰) in higher trophic level species, due to the greater dietary protein quality of higher trophic level predators (Robbins et al., 2005). Since discrimination factors for bathyal megafauna are yet to be determined, we used discrimination factors of 3.15 ± 1.28‰ for δ^15^N and 0.97 ± 1.08‰ for δ^13^C (Sweeting et al., 2007), which have been previously used to study the trophic structure of meso- and bathypelagic fishes in the Gulf of Mexico (Richards et al., 2018) and in the Western Mediterranean Sea (Valls et al., 2014). Each model was run with identical parameters (number of MCMC chains = 3; chain length = 300000; burn in = 200000; thin = 100), and model convergence was determined using Gelman-Rubin and Geweke diagnostic tests (Stock et al., 2018).

## 3 Results

### 3.1 Stable isotopes

16.14‰ for crustaceans. Fish mean δ^13^C values differed by 4.27‰, separating the most depleted (*Macrorhamphosus scolopax:* −20.85 ± 0.46‰, sampling depth of 200 m) and the most enriched species (*Centrophorus granulosus* and *Conger conger*: −16.89 and −16.57‰, respectively, sampling depth of ~1000m) (**Table 1**, **Figure 2**). Crustaceans species-specific mean δ^13^C varied less by 2.07‰, where the most depleted species was *Parapenaeus longirostris* (−18.45 ± 0.10‰, sampling depth of 200 m) and the most enriched species was *Polycheles typhlops* (−16.38 ± 0.21‰, sampling depth of ~1400 m). Species-specific differences in δ^13^C and δ^15^N were significant for both fish (MANOVA, *F*_13,144_ = 19.73, p < 0.001) and crustaceans (MANOVA, *F*_5,78_ = 14.62, p < 0.001).

**Figure 2.**
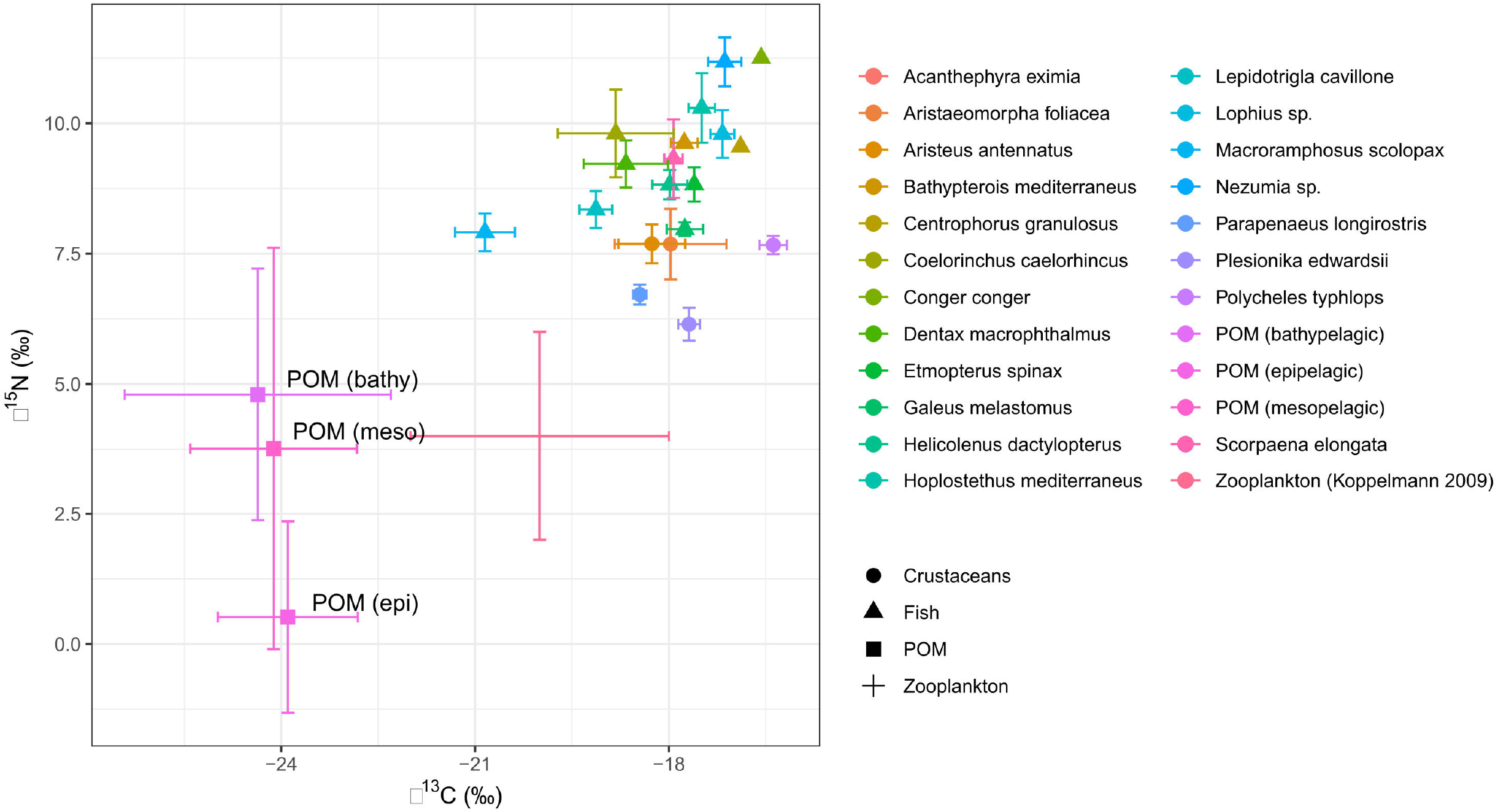
Isotope biplot of δ^13^C and δ^15^N values of Levantine deep-sea decapod crustaceans (circles) and fishes (triangles), zooplankton (cross, based on Koppelmann et al. 2009), and POM (squares). Data points represent means and error bars represent ±SD.

Species-specific mean δ^15^N values varied from 11.36 ± 0.39‰ (*Nezumia* sp. 1100 m depth) to 7.91 ± 0.36‰ (*M. scolopax*, 200 m depth) in fish and from 6.15 ± 0.31‰ (*Plesionika edwardsii;* 200-600 m depth) and 8.07 ± 0.21‰ (*Aristaeomorpha foliacea*; 1400 m depth) in crustaceans. Fish mean δ^15^N values positively correlated with the δ^13^C values (*r*^2^ = 0.6, p < 0.001, **Figure 2**, **Supplementary Figure 1**) and varied among species (ANOVA, *F*_13,72_ = 24.22, p < 0.001). Crustaceans, however, did not show this correlation between δ^15^N and δ^13^C (*r*^2^ = 0.002, p > 0.05, **Figure 2**, **Supplementary Figure 2**), observed in fish from similar depths. Due to limited spatial coverage within each species, spatial variation could not be tested within each species, and therefore, spatial trends were tested by addressing all fish species together. Fish δ^13^C values positively varied with bottom depth (*r*^2^ = 0.42; P <0.01, **Figure 3**), where the most enriched samples are found at the continental rise (> 1000 m) and the most depleted at the shallow slope (200m) at the edge of the shelf. This pattern was less clear in the case of fish δ^15^N values (**Figure 4**), where species-specific mean values seem more variable in the continental rise (> 1000 m). Crustaceans mean δ^15^N values positively correlated with depth (*r*^2^ = 0.76, p = 0.053, **Figure 4**), while their mean δ^13^C values showed no such correlation (**Figure 3**).

**Figure 3.**
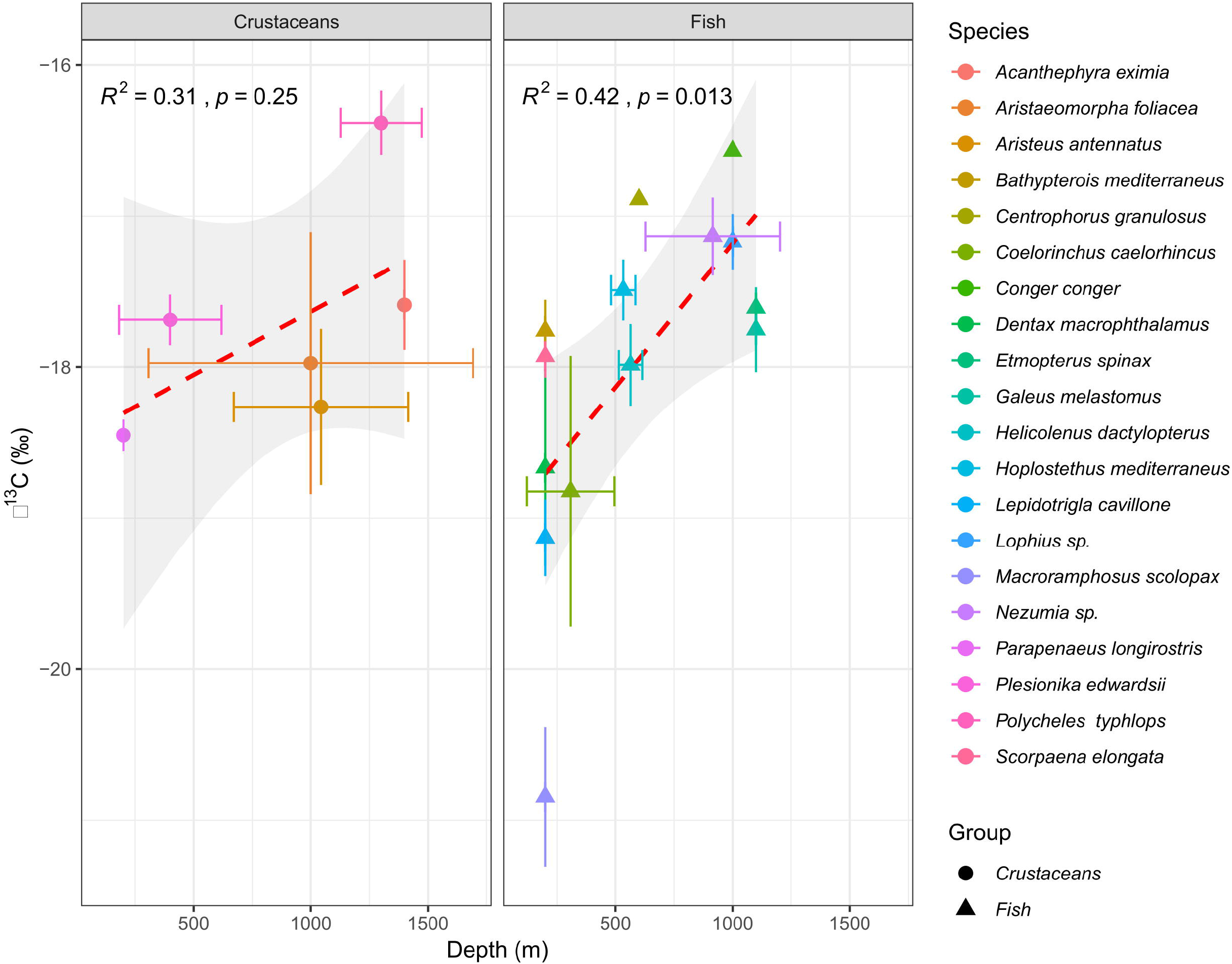
Correlations between mean δ^13^C (‰) and depth (m) in Levantine deep-sea decapod crustaceans (left) and fish (right). Dash lines represent Least-squares regression line.

**Figure 4.**
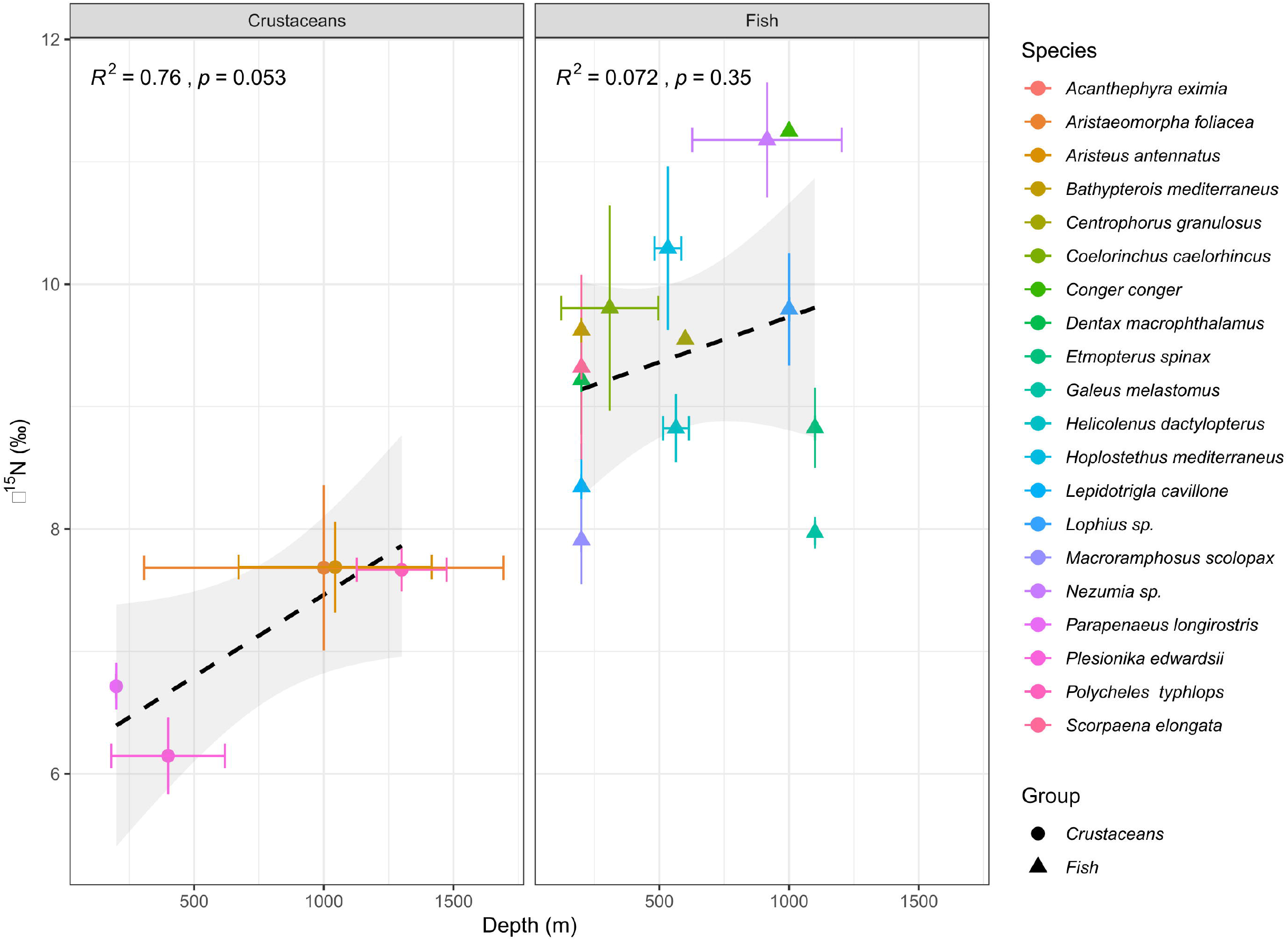
Correlations between mean δ^15^N (‰) and depth (m) in Levantine deep-sea decapod crustaceans (left) and fish (right). Dash lines represent Least-squares regression line.

POM collected from depths ranging from 0 to 1135 m, exhibited a wide δ^13^C range (−27.36 to – 21.54‰) and δ^15^N range (−3.25 to 12.76‰), with POM samples generally becoming more enriched in ^15^N and more depleted in ^13^C at bottom depths (**Figure 5**). Significant differences in POM δ^13^C and δ^15^N among the three vertical depth zones were observed (MANOVA, *F*_2,148_ = 8.65, p < 0.001). POM-δ^13^C and C/N ratio exhibited a negative correlation (**Supplementary Figure 3**), which was not observed in POM-δ^15^N and C/N ratio.

**Figure 5.**
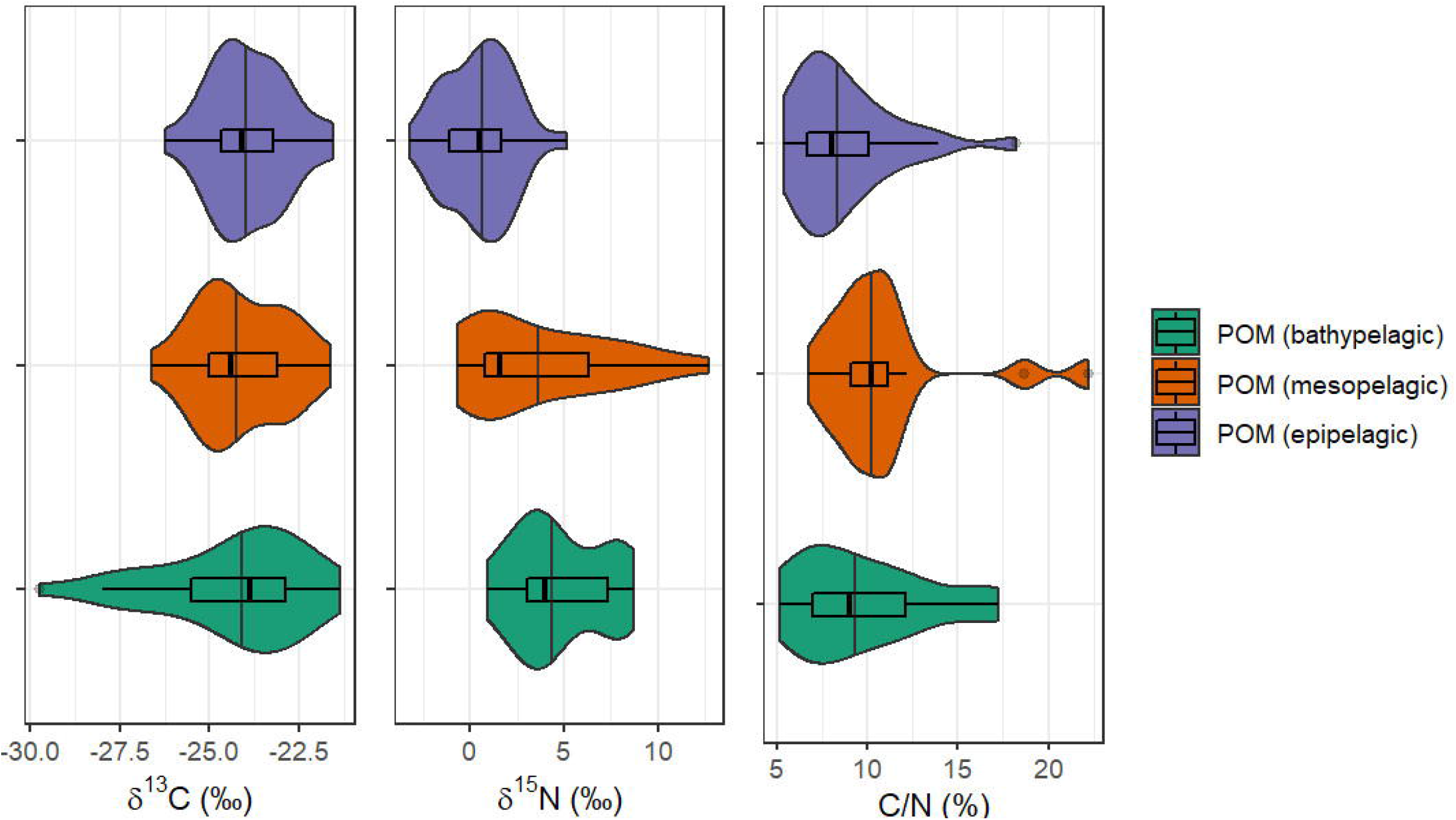
Violin charts with boxplots showing POM-δ^15^N (‰), POM-δ^13^C (‰), and POM-C/N ratio collected from epipelagic (0-200 m), mesopelagic (201-800 m), and bathypelagic (>800 m) depths in the Southeastern Mediterranean Sea during 2018-2021.

### 3.2 Trophic level estimates

We used the average epipelagic POM-δ^15^N value (0.52 ± 1.84‰) as a baseline for estimating species-specific TL, following Richards et al. (2018). The δ^15^N of bathypelagic primary consumers was not available for similar calculation. However, based on the low δ^15^N values of zooplankton in the eastern Mediterranean, it was assumed that the primary food source, namely smaller zooplankton, phytoplankton and particles has a δ^15^N value around zero (Koppelmann et al., 2009). Large mesozooplankton (333 mm mesh size, upper water column) δ^15^N values in the EMS show an enrichment trend across a west-east transect (SE Crete mean δ^15^N value ~2.0‰ and SE Cyprus mean δ^15^N value ~4.0‰, Koppelmann et al. 2009). The average Δ^15^N of phytoplankton-zooplankton and zooplankton-fish in the study area yielded enrichment factors of 2.5 and 3.9‰, respectively (Grossowicz et al. 2019). Using this factor, our measured epipelagic POM-δ^15^N value (0.52 ± 1.84‰) could be also inferred and confirmed using the large mesozooplankton δ^15^N value of ~4.0‰ reported by Koppelmann et al. (2009). When epipelagic POM data were used to set the baseline, fish TL ranged from 3.85 ± 0.11 (*M. scolopax;* 200 m depth) to 4.91 (*C. conger*; 1000 m depth), while the average TL of all fish species was 4.33 ± 0.34 (**Figure 6**). Crustaceans-δ^15^N values yielded TLs between 3.29 and 3.78, with an average of 3.62 ± 0.22 (**Figure 6**).

**Figure 6.**
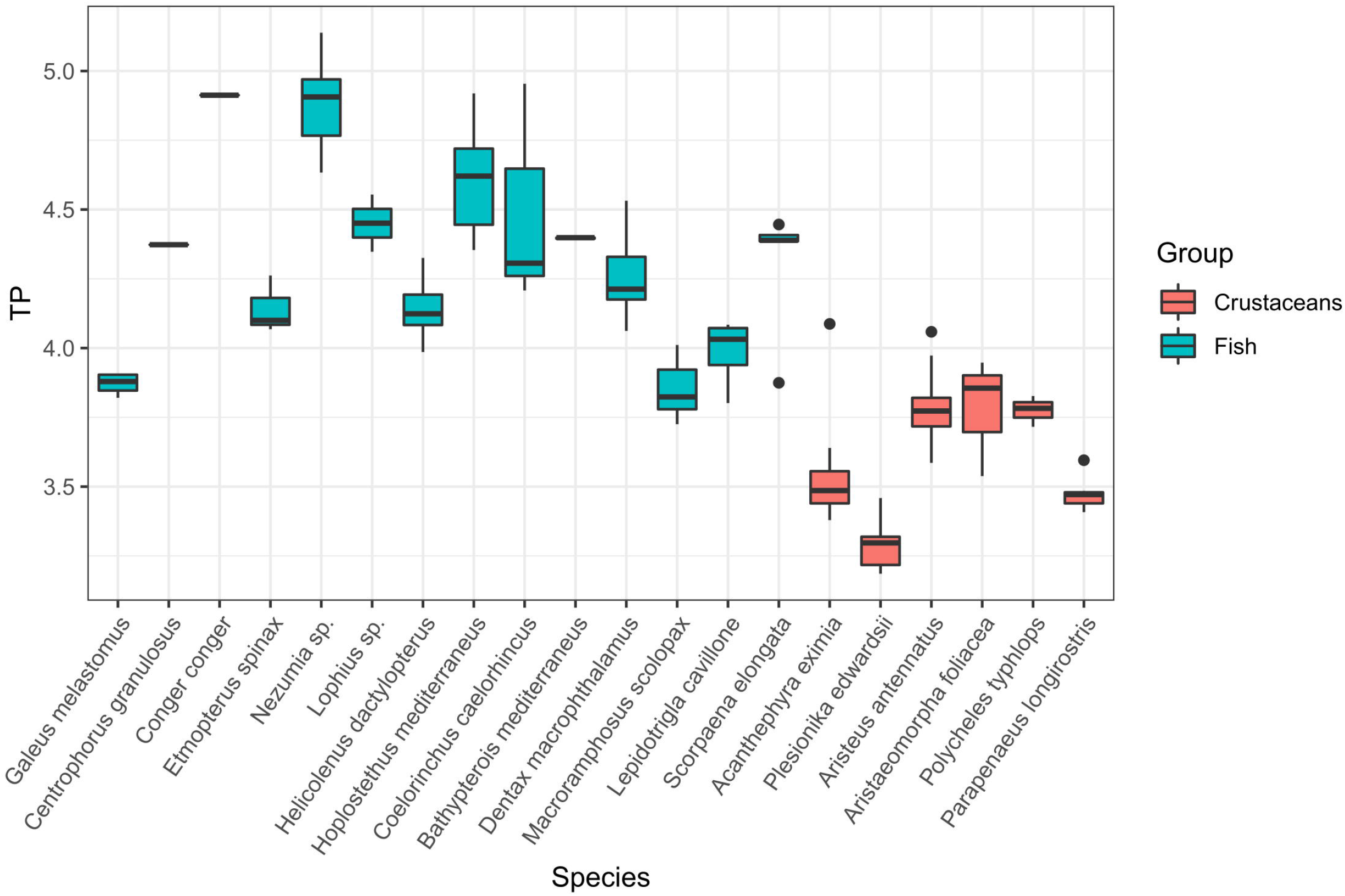
Trophic level (TL) estimates calculated using δ^15^N data of deep-sea Levantine decapod crustaceans (red) and fish (blue) species, using epipelagic POM to establish isotopic baseline.

Of the species examined, only few enabled an estimation of ontogenetic effect (**Figure 7**). This is due to the low range of body size within individual species that were sampled in this study. Nevertheless, the crustaceans *A. eximia* (*r*^2^ = 0.82; p < 0.001) and *P. edwardsii* (*r*^2^ = 0.72; p < 0.05) exhibited a positive relationship between length and δ^15^N values. Size and δ^13^C values did not yield significant correlations. Positive relationship between size and δ^15^N values was also observed for the fish *Dentex macrophthalmus* (*r*^2^ = 0.59; p < 0.05).

**Figure 7.**
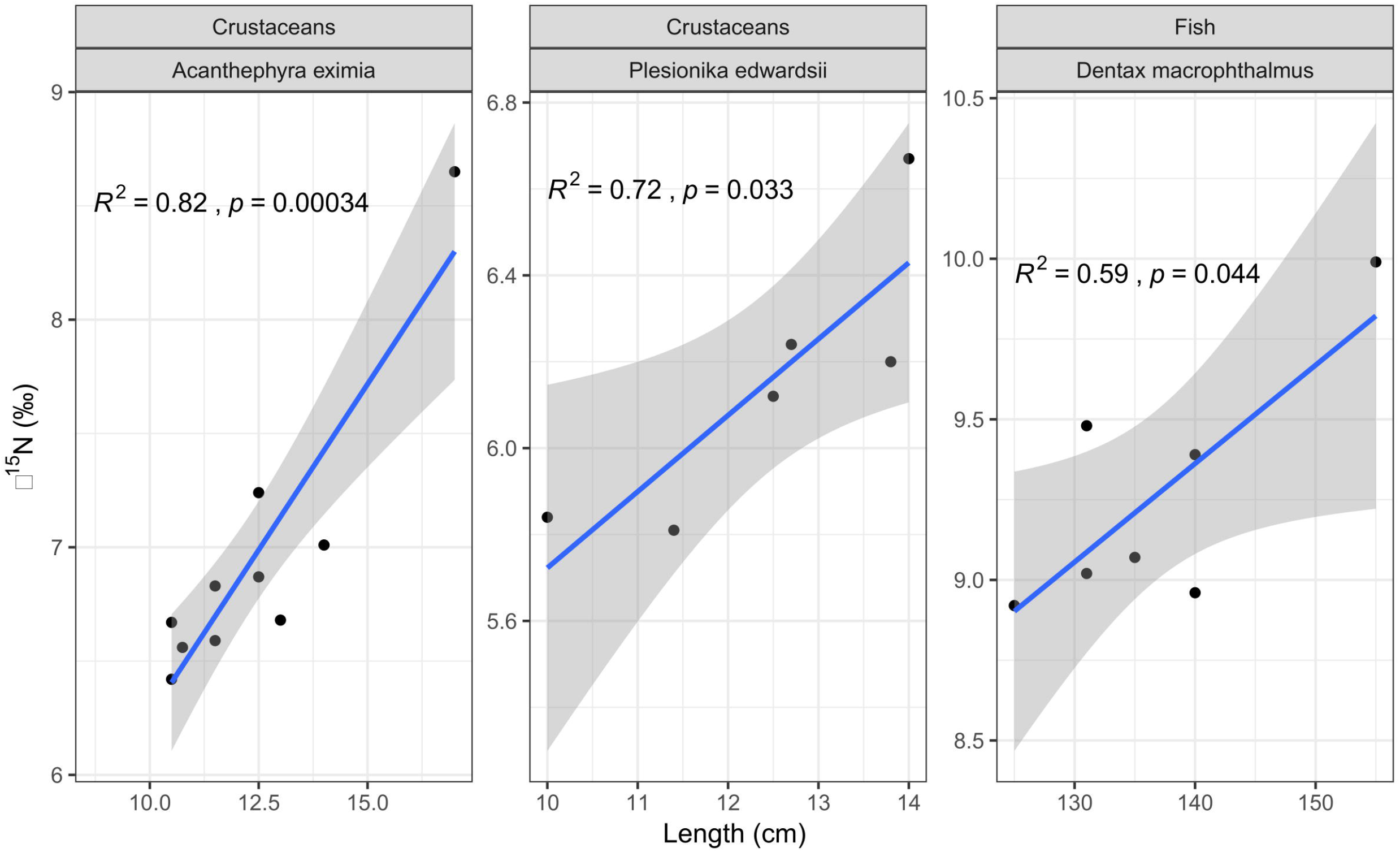
Least-squares regression analysis between total length (cm) and δ^15^N values in the decapod crustaceans *A. eximia* (*n*=10) and *P. edwardsii* (*n*=6), and the fish *D. macrophthalmus* (*n*=7).

### 3.3 Trophic niche breadth

Isotopic niche breadth, calculated using SEAc (**Table 2**, **Figure 8**), was largest for the fish collected from the shallow continental slope *Coelorinchus caelorhincus* (SEAc = 2.04), *D. macrophthalmus* (SEAc = 0.94) and *M. scolopax* (SEAc = 0.66), and for the shrimps *A. foliacea* (SEAc = 0.96) and *Aristeus antennatus* (SEAc = 0.63), both opportunistic carnivores. The smallest isotopic niche breadth belonged to *Bathypterois mediterraneus* (SEAc = 0.009), a planktivorous fish. Fish and crustacean community metrics (**Table 3**) showed higher convex hull area (TA) in fish (TA = 7.37) than in crustaceans (TA = 0.81), indicating a larger trophic community width (Layman et al. 2007).

**Table 2:** Metrics for estimating isotopic niche size in fourteen fish and six decapod crustaceans from the Levantine bathyal (*n*≥3). TA, total area (‰^2^) encompassed by all data points of each species; SEA, standardized ellipse area for each species; SEA_C_, size-corrected standardized ellipse area.

| Sample type / Species | TA | SEA | SEAC |
| --- | --- | --- | --- |
| **Fish** |  |  |  |
| *B. mediterraneus* | 0.00 | 0.00 | 0.01 |
| *C. caelorhincus* | 3.20 | 1.84 | 2.04 |
| *D. macrophthalmus* | 1.61 | 0.85 | 0.94 |
| *E. spinax* | 0.01 | 0.03 | 0.05 |
| *G. melastomus* | 0.09 | 0.10 | 0.15 |
| *H. dactylopterus* | 0.53 | 0.21 | 0.22 |
| *H. mediterraneus* | 0.34 | 0.25 | 0.31 |
| *L. cavillone* | 0.31 | 0.27 | 0.34 |
| *M. scolopax* | 0.60 | 0.49 | 0.66 |
| *Nezumia* sp. | 0.90 | 0.37 | 0.41 |
| *S. elongata* | 0.24 | 0.24 | 0.32 |
| **Crustaceans** |  |  |  |
| *A. eximia* | 0.90 | 0.50 | 0.56 |
| *A. foliacea* | 0.27 | 0.48 | 0.96 |
| *A. antennatus* | 1.91 | 0.59 | 0.63 |
| *P. longirostris* | 0.08 | 0.06 | 0.07 |
| *P. edwardsii* | 0.21 | 0.16 | 0.20 |
| *P. typhlops* | 0.03 | 0.06 | 0.12 |

**Figure 8.**
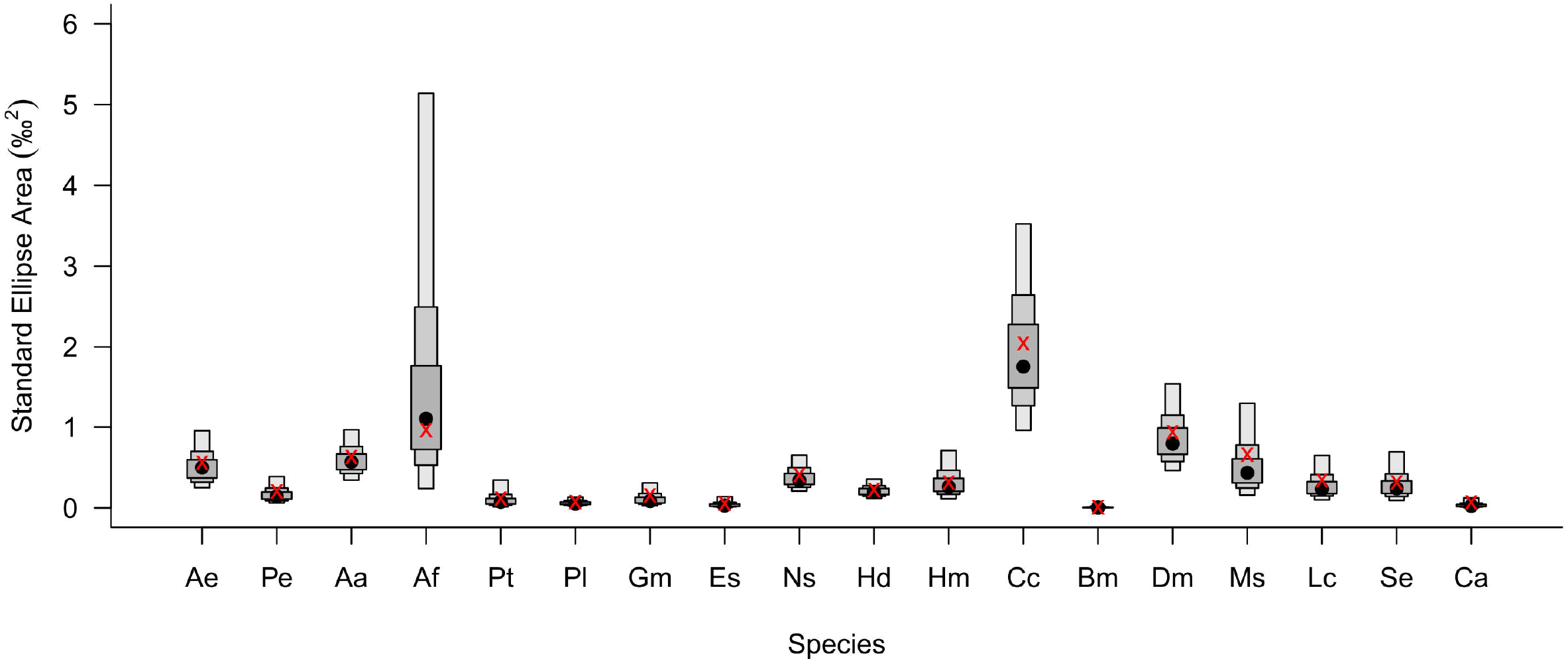
Trophic niche breadth of Levantine deep-sea fish and decapod crustaceans (*n*≥3) estimated by size-corrected standard ellipse area (SEA_B_) boxplots (showing 95, 75 and 50 % credibility intervals). Black circles represent means; red x symbols represent the maximum likelihood estimates of SEA_B_. Ae - *Acanthephyra eximia*; Pe - *Plesionika edwardsii*; Aa - *Aristeus antennatus*; Af - *Aristaeomorpha foliacea*; Pt - *Parapenaeus longirostris*; Pl - *Parapenaeus longirostris*; Gm - *Galeus melastomus*; Es - *Etmopterus spinax*; Ns - *Nezumia* sp.; Hd - *Helicolenus dactylopterus*; Hm - *Hoplostethus mediterraneus*; Cc - *Coelorinchus caelorhincus*; Bm - *Bathypterois mediterraneus*; Dm - *Dentex macrophthalmus*; Ms - *Macroramphosus scolopax*; Lc - *Lepidotrigla cavillone*; Se - *Scorpaena elongata*.

**Table 3:** Community metrics for estimating isotopic niche size (Layman et al. 2007) in bathypelagic fish and bathybenthic crustaceans. TA, total area (‰^2^) encompassed by all data points of each assemblage (crustaceans, fish); CD, mean distance to centroid (trophic diversity); MNND, mean nearest neighbour distance (trophic similarity); SDMNND, the standard deviation of MNND (trophic evenness).

|  | Crustaceans | Fish |
| --- | --- | --- |
| $\delta^{13}\text{C}$ range | 1.54 | 4.67 |
| $\delta^{15}\text{N}$ range | 2.07 | 3.71 |
| TA | 1.87 | 7.37 |
| CD | 0.81 | 1.29 |
| MNND | 0.75 | 0.82 |
| SDNND | 0.43 | 0.52 |

### 3.4 Bayesian mixing models

The results of the mixing models indicate that most deep-sea fish and crustacean consumers included in this study derive the bulk of their carbon from bathypelagic POM (**Figure 9**). Relative contributions of epi- and mesopelagic POM ranged from 0.3 ± 0.3% in *Nezumia* sp. to 17.1 ± 8.1% in *Lepidotrigla cavillone*, while contributions from bathypelagic POM were much higher, ranging from 73.7 ± 4.3% in *L. cavillone* to 99.3 ± 0.5% in *Nezumia* sp., with the exception of *M. scolopax* that had similar bathypelagic and mesopelagic contribution (45.7 ±5.0% and 40.4 ± 12.4%, respectively). Diagnostic plots of posterior distributions (**Supplementary Figure 4**) revealed high negative correlations between bathypelagic POM and epi/mesopelagic POM (*R_epi-bathy_* = −0.78, *R_meso-bathy_* = −0.49) and a low negative correlation between epi- and mesopelagic POM (*R_epi-meso_* = −0.17). The negative correlation is likely caused by the similar δ^13^C signatures of sources and not from a missing carbon source, since producer data fully constrain consumer data when an appropriate trophic enrichment factor is applied (Richards et al., 2018) and that model diagnostics indicate that the model fully converged (Gelman-Rubin Diagnostic: all variables <1.01; Gweke Diagnostic: <5% of variables outside ± 1.96 for each chain).

**Figure 9.**
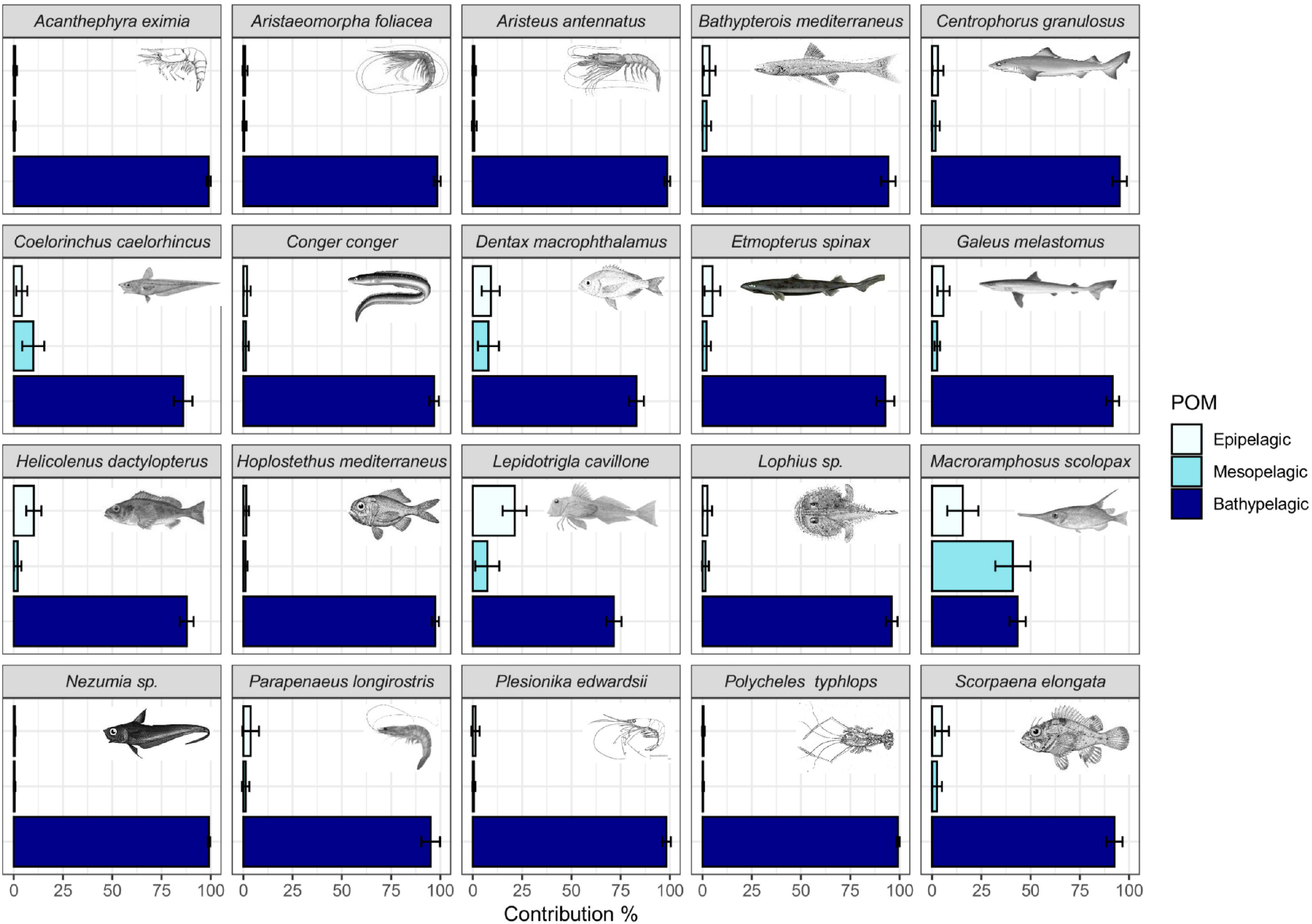
Estimated relative contributions (%) of POM collected from epi-, meso- and bathypelagic depths (0-200, 201-800, >800 m, respectively) to Levantine deep-sea fish and decapod crustaceans, based on Bayesian mixing models. Bars represent mean contributions and error bars represent ±SD.

## 4 Discussion

This is the first attempt to elucidate the trophic ecology of deep-sea fish and crustacean species in the SEMS. The knowledge gained in this study provides insights into the main energy sources sustaining deep-sea food webs in one of the most oligotrophic, nutrient-improvised marine basins, worldwide. However, insights gained in this study are not limited to the SEMS alone, and can be ascribed to many oligotrophic basins with limited carbon and nutrient sources.

Our δ^13^C and δ^15^N values varied across fish species and as a function of bathymetric depth, suggesting that depth and diet are controlling the trophic positions inferred from our stable isotope data. As expected, top predators such as the European conger eel *C. conger*, occupied the highest trophic position. The rattail *Nezumia* sp., a small macrourid fish that was collected from similar depths of >1000 m, yielded similar high δ^15^N values. Both species occupied a maximum trophic level of 4.89-4.91. Polunin et al. (2001) found similar trophic position of 4.4 for both the shark *Centroscymnus coelolepsis* and *Nezumia aequalis* in the continental slope of the Balearic Islands. High δ^15^N values of *Nezumia* (11.09 ±0.58‰ and 11.31‰) were also recorded by Fanelli and Cartes (2010) in the Archipelago of Cabrera (Algerian Basin) and by Papiol et al. (2013) in the Balearic Islands (Catalan Sea, West Mediterranean), respectively, and were attributed to the suprabenthic crustaceans and polychaetes that constitute the diet of this macrourid. Our TL data also agree well with that of benthic carnivorous fish from Bay of Banyuls-sur-Mer (northwest Mediterranean, France; Carlier et al., 2007). Among the fish, the lowest trophic level (3.85±0.11) was found in the snipefish *M. scolopax*, which feeds on hyperbenthic demersal zooplankton during daytime (Carpentieri et al., 2016). This was also inferred from the results of the mixing models, indicating a relatively high contribution of mesopelagic POM to the diet of *M. scolopax*. Relatively to the fish, the bathybenthic crustaceans measured in this study occupied lower trophic positions – between 3.29 and 3.78, in agreement with the TL of deep benthic invertebrates of the Western Mediterranean (Carlier et al., 2007).

In the fish species examined here, mean δ^15^N values spanned 3.45‰, about 1.1 TL, while in the crustacean species mean δ^15^N values spanned 1.92‰, about 0.6 TL (assuming trophic enrichment factor of 3.15‰). Our observed ranges of estimated trophic levels are in line with other studies examining Mediterranean (1.1 TL, Valls et al., 2014), Pacific (1.6 TL, Choy et al., 2015), and the Gulf of Mexico (0.62 TL, Richards et al., 2018). Different feeding strategies as well as different migration habits may explain wider range of δ^15^N (Shipley et al., 2017a;Richards et al., 2020). Despite of the reliance on similar basal production, mesopelagic fishes from the Western Mediterranean were segregated by trophic position, between 2.9 for the small bristlemouth *Cyclothone braueri* to 4.0 for the lanternfish *Lobianchia dofleini* (Valls et al., 2014), and bathyal fishes off the Balearic Islands appeared to be foraging over two to three full trophic levels (Polunin et al., 2001). Our results support a much narrower trophic range for bathyal fish and bathybenthic decapod crustaceans in the SEMS. We attribute this narrow range to the ultra-oligotrophic state of the SEMS, resulting in limited carbon sources to sustain the deep-sea food webs, reflected by a general increase of δ^13^C in fish as function of bottom depth. This pattern could be driven by a number of factors including shifting production sources, or shifts in community composition and feeding strategies, and or switching from benthic to pelagic prey (Fanelli et al., 2011;Trueman et al., 2014). For example, ^13^C became more depleted in individuals captured at greater depths in the deep-sea island slope system of the Exuma Sound, the Bahamas (Shipley et al., 2017b). Inshore-to-offshore depletion in ^13^C values were also apparent in epipelagic fishes in the northern California Current, where copepods, gelatinous zooplankton, and nekton showed a significant linear decrease in δ^13^C with distance offshore (Miller et al., 2008).

The major carbon sources supporting deep-sea food webs are poorly defined, aside from oligotrophic open-ocean gyres, where sinking phytoplanktonic-POM is considered the main energy source (Shipley et al., 2017b). This was observed by a narrow range of δ^13^C in meso- and bathypelagic predatory fishes in the Gulf of Mexico, indicating similar epipelagic carbon source (Richards et al., 2018). Conversely, the results of our mixing-models show that the majority of carbon (92.19 ± 12.54%) supporting the species examined in this study is not derived from epipelagic sources. An alternative hypothesis is that the source of carbon in the deep-sea originates from the shelf. A significant proportion of neritic-derived primary production may be transported into deep-sea systems by currents (Suchanek et al., 1985;Sanchez-Vidal et al., 2012;Efrati et al., 2013), or via lateral transport (Fahl and Nöthig, 2007), and once assimilated into the food web, more enriched ^13^C values are to be expected (Polunin et al., 2001;Fanelli et al., 2011). Katz et al. (2020) used deep-sea sediment traps in the Israeli Southeastern Mediterranean Sea and showed that lateral transport from the continental margin contributes the greatest fraction of particulate flux to the seafloor. Therefore, we suggest that lateral transport constitutes the main source of carbon to the deep-sea food web in the Southeastern Mediterranean Sea.

Since the carbon signature of primary producers can significantly vary between macroalgae and different phytoplankton groups (Fanelli et al., 2011;Grossowicz et al., 2019), food webs that show a linear relationship between δ^15^N and δ^13^C values are suggestive of a single food source (Polunin et al., 2001;Carlier et al., 2007). Generally weak δ^13^C–δ^15^N correlations were found in deep-sea macrozooplankton and micronekton off the Catalan slope likely due to the consumption of different kinds of sinking particles (e.g. marine snow, phytodetritus). Multiple recycling of POM constituted an enrichment effect on the δ^13^C and δ^15^N values of deep-sea macrozooplankton and micronekton (Fanelli et al., 2011). Our results yielded significant positive correlation between fish δ^15^N and δ^13^C values, further supporting a single food source.

Our results showed that δ^13^C fractionates less than 1.0‰ for each trophic position. The Δ^13^C between the mean water column POM-δ^13^C (−24.13 ± 1.56‰) and fish/crustaceans δ^13^C (−18.10 ± 0.93‰) of the SEMS amounted to 6.04‰ (equal to at least six trophic positions), and therefore, cannot be attributed to trophic enrichment, but rather to the regeneration of benthic carbon sources. Moreover, our δ^13^C-C/N data support the potential effect of microbially degraded phyto-detritus resulting in higher isotopic values of nitrogen and carbon in deep benthic food webs compared with pelagic food web (Papiol et al., 2013;Romero-Romero et al., 2021).

Deep-sea ecosystems are subjected to exacerbating anthropogenic stressors, including overfishing, chemical pollution, mining, dumping, litter, plastics, and climate change (Davies et al., 2007). In oligotrophic environments such as the ultra-oligotrophic SEMS, deep-sea ecosystems are further vulnerable to reduced food availability (Kröncke et al., 2003). Regeneration of benthic carbon sources, supported by this study, provides oligotrophic deep-sea food webs with a greater ability to endure carbon limitation. Nonetheless, benthic carbon source originating in lateral transport from the shallow shelf to the deep-sea, as indicated here, may carry detrimental implications to the ecosystem via pollutant accumulation and biomagnification (Liu et al., 2020). This is particularly important in marginal seas that are prone to anthropogenic pollution (Kim et al., 2019;Shoham-Frider et al., 2020). Continuous studies should be undertaken to further unveil the implications of lateral transport and benthic carbon regeneration to deep-sea food webs.

## Supporting information

Trophic ecology of deep sea ecosystem bioRxiv Supplementary

## 4 Data Availability Statement

The datasets generated for this study can be found in the open-access data repository PANGAEA, and will be made publicly available with publication.

## 8 Author Contributions

G.S-V. and T. G-H. conceived this study. G.S-V., N.S. and T.G-H. collected the data. N.S. provided species identification and measurements. G.S-V. and T. G-H. analyzed and modeled the stable isotope data. All coauthors contributed substantially to drafting the manuscript, and approved the final submitted manuscript.

## 9 Acknowledgments

The authors wish to thank Mor Kanari for map preparation, and the captain and crew of R/V Bat-Galim, including onboard technical and scientific personnel for assistance in sampling. This work was supported by the National Monitoring Program of the Israeli Mediterranean waters.

