## Supplementary material for "Trophic ecology of deep-sea megafauna in the ultra-oligotrophic Southeastern Mediterranean Sea": Trophic ecology of deep sea ecosystem bioRxiv Supplementary

### 1 Supplementary Figures and Tables

#### 1.1 Supplementary Figures

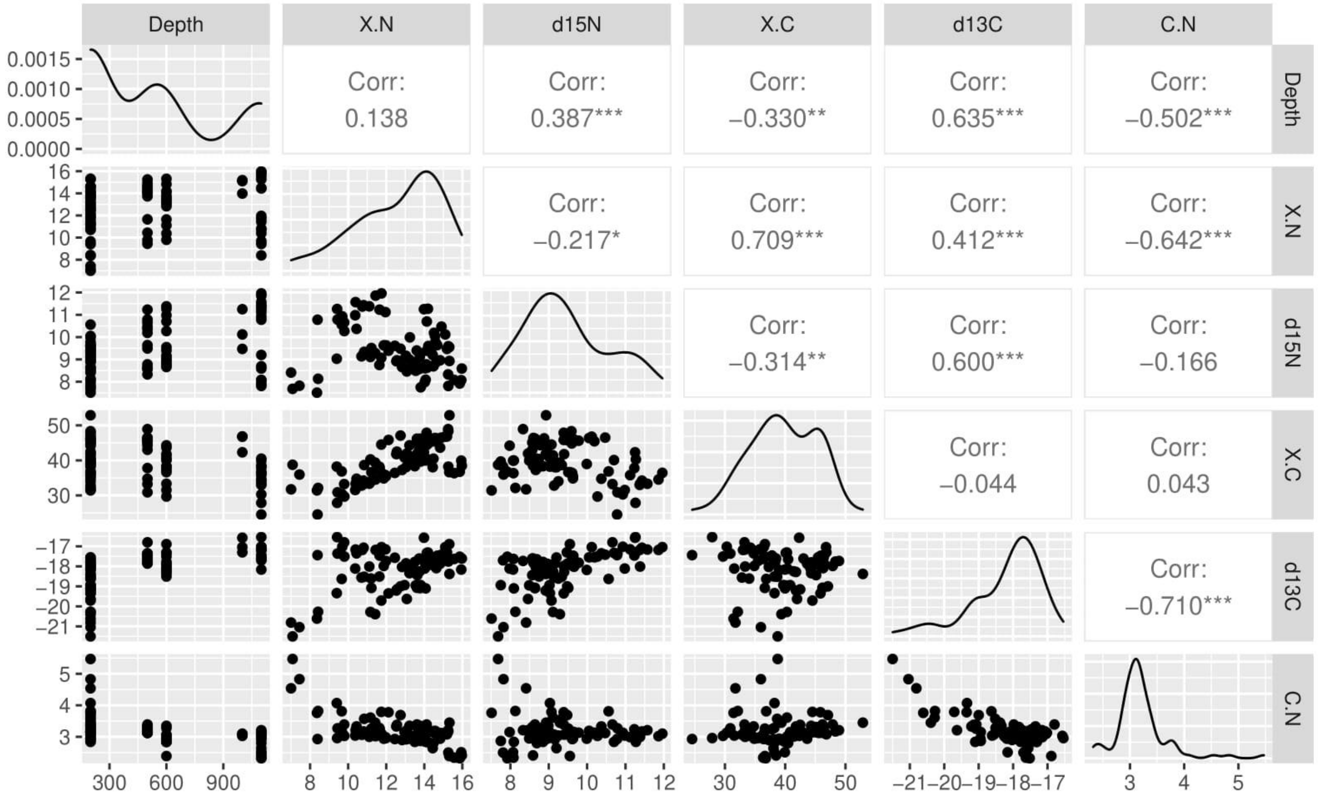


**Supplementary Figure 1**. Pairwise correlation matrix between depth, %C, $\delta^{13}$C, %N, $\delta^{15}$N and C/N ratio in bathypelagic fish (n=86) collected from the Southeastern Mediterranean Sea. Data points are shown in the scatterplots below the diagonal. Variable distributions are presented on the diagonal. Pearson correlation coefficients are presented in the cells above the diagonal, asterisk denotes significance level.

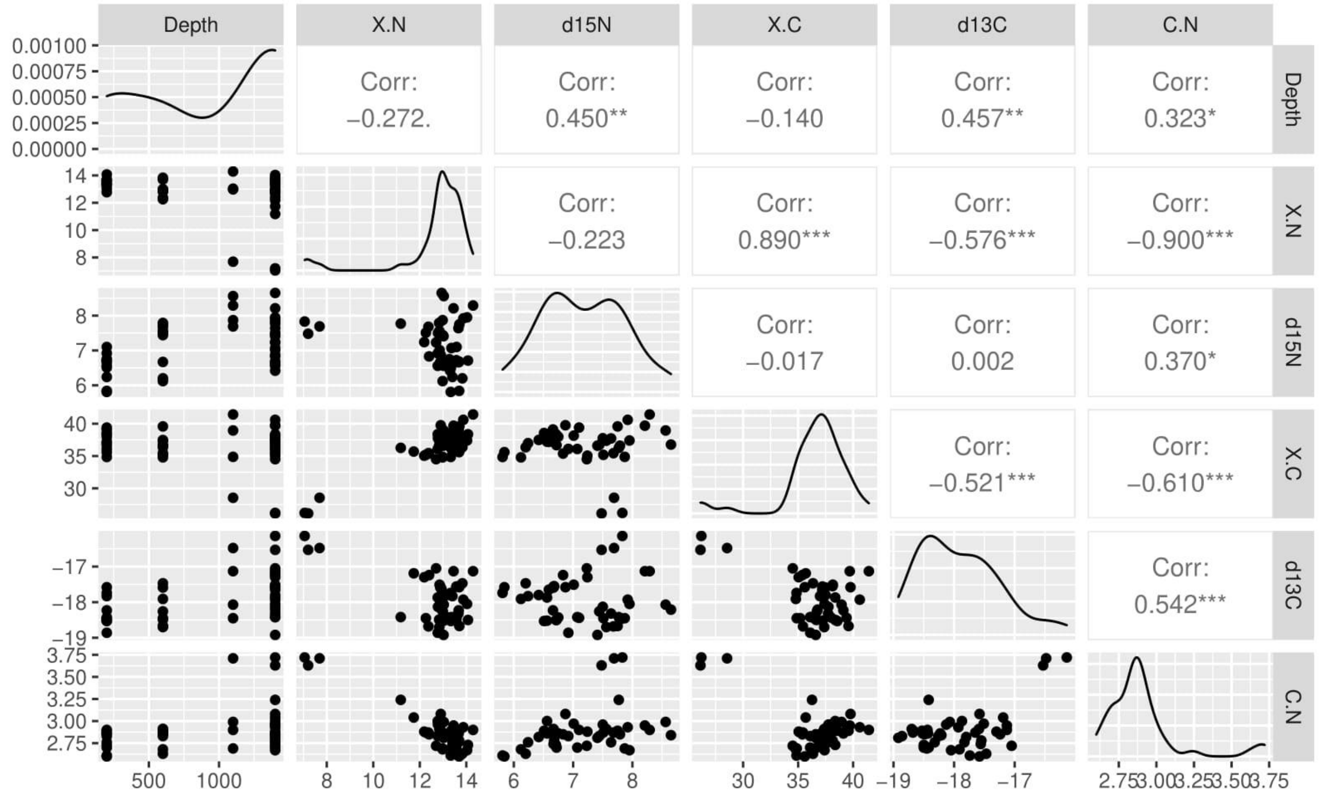


**Supplementary Figure 2**. Pairwise correlation matrix between depth, %C, $\delta^{13}$C, %N, $\delta^{15}$N and C/N ratio in bathybenthic crustaceans (n=46) collected from the Southeastern Mediterranean Sea. Data points are shown in the scatterplots below the diagonal. Variable distributions are presented on the diagonal. Pearson correlation coefficients are presented in the cells above the diagonal, asterisk denotes significance level.

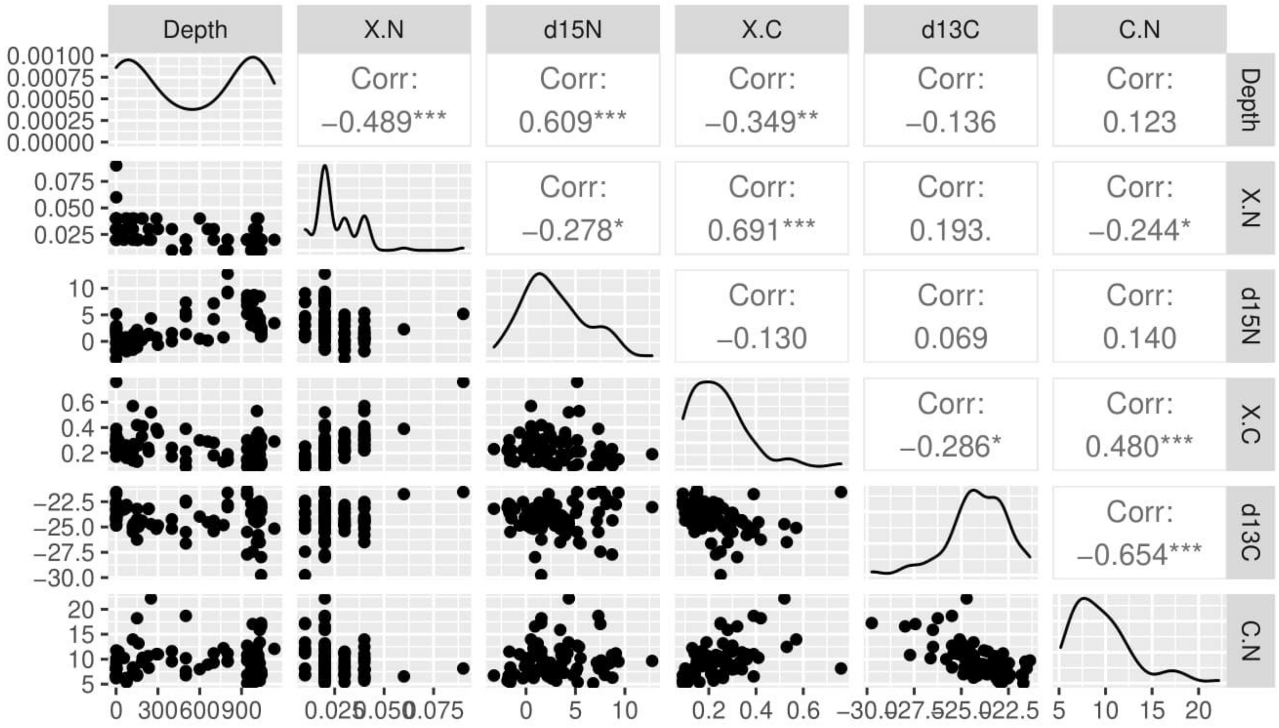


**Supplementary Figure 3**. Pairwise correlation matrix between depth, %C, $\delta^{13}C$, %N, $\delta^{15}N$ and C/N ratio in POM samples (n=77) collected from the Southeastern Mediterranean Sea. Data points are shown in the scatterplots below the diagonal. Variable distributions are presented on the diagonal. Pearson correlation coefficients are presented in the cells above the diagonal, asterisk denotes significance level.

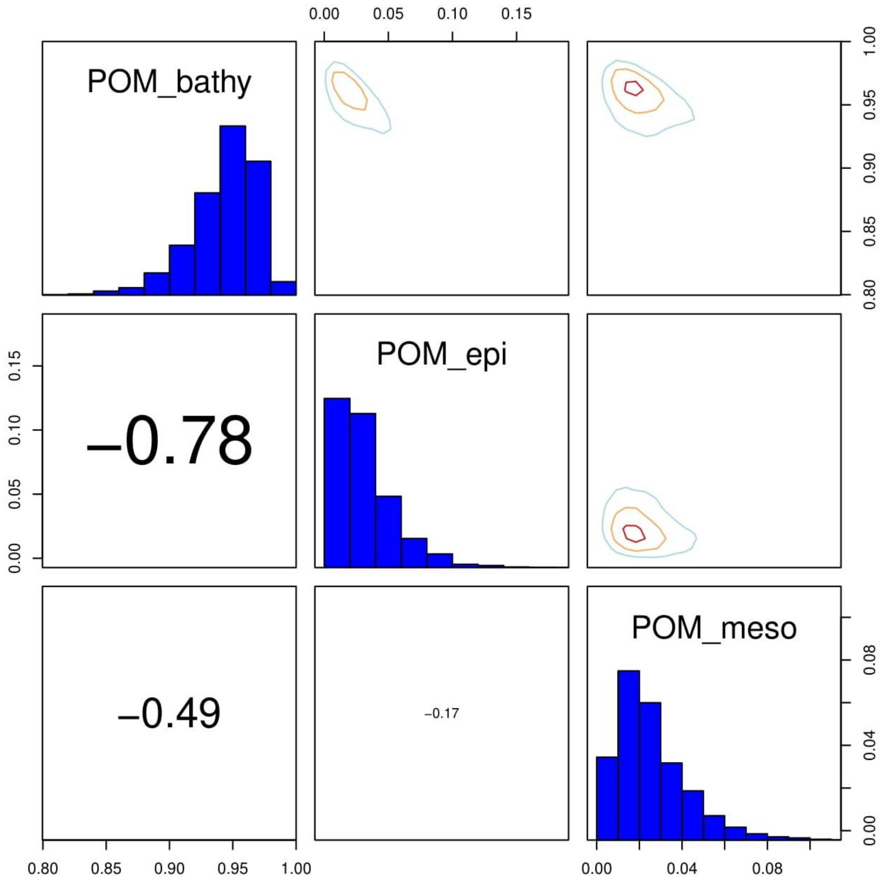


**Supplementary Figure 4**. Matrix plot presenting estimated POM type (epipelagic, mesopelagic, bathypelagic) proportions contributing to Levantine bathyal megafauna calculated using Bayesian mixing models. The diagonal cells show the posterior probability distributions for each of the POM types. The cells below the diagonal show the correlations between contributions for pairs of POM types. The cells above the diagonal show contours of the joint posterior probability distribution for contributions for pairs of POM sources.
